# Neurovascular unit adjustments following chronic distress explain motivational deficits in mice

**DOI:** 10.1101/2024.03.25.586557

**Authors:** Lidia Cabeza, Damien Mor, Bahrie Ramadan, Guillaume Benhora-Chabeaux, Christophe Houdayer, Emmanuel Haffen, Yvan Peterschmitt, Adeline Etievant, Fanchon Bourasset

## Abstract

**Background:** The neurovascular unit (NVU) represents the structural and functional relationship between the neural tissue and the blood. Neurovascular dysfunction has been highlighted in neuropsychiatric afflictions, but whether it is a cause or a consequence of the pathology remains to be elucidated. Thus, to elucidate the role of the NVU on the emergence of emotional-cognitive dysfunction, it is necessary to study how its individual components associate. This study therefore aims at investigating whether the development of depressive-related loss of motivation is grounded on NVU adjustments impacting the permeability of the blood-brain barrier (BBB) and in particular, of the structural scaffolding of microvessels.

**Methods:** Adult male C57BL/6jRj mice chronically treated with corticosterone (CORT) and showing severe motivational deficits in an operant progressive ratio (PR) schedule of reinforcement task, presented altered neural activation assessed through FosB expression in key brain regions involved in motivational processing (anterior insular cortex, basolateral amygdala, bed nucleus of the stria terminalis and ventral tegmental area). We evaluated NVU modifications through immunofluorescence staining targeting specific markers of microglia (IBA-1), endothelial tight junctions (ZO-1) and astrocytes (GFAP). The effect of chronic CORT administration on mice BBB permeability was evaluated through *in vivo* perfusion of fluorescent 40 kDa Dextran.

**Results:** Our results highlight that where sustained neuronal activation failed, NVU modifications predict behavioural deficits in CORT-treated animals. Notably, our analyses show that NVU modifications within the ventral tegmental area are essential to understand effort-based related behavioural performance in mice, and most particularly, that the key element of microvessels’ tight junctions ZO-1 plays a pivotal role on motivation-related behavioural output.

**Conclusions:** Our results confirm a direct role of neurovascular adaptations on emotional and cognitive behavioural performance in mice, and therefore place the NVU in a key position in the research of the biological substrate at the origin of neuropsychiatric disorders.

## Background

Prolonged stress exposure deregulates stress-related adaptive mechanisms and cognitive function, which often lead to neuropsychiatric disorders such as anxiety and major depression (1,2). An estimated 3.8% of the world population, i.e. 280 million people, experience depression, including mild and severe episodes (3).

Among the core symptoms defining major depression, anhedonia or loss of pleasure or interest for common activities and altered self-motivation significantly affect individuals’ initiative and decision-making capability. Motivation is therefore considered a key modulator of reward-related behaviour, which impacts high-order functioning by influencing an individual’s engagement in rewarding behaviours, from initiation (appetitive behaviour) to accomplishment (consummatory behaviour) by maintenance through persistent effort (4–6).

Previous works from our laboratory have consistently demonstrated that chronic-distress related human pathology can be successfully reproduced in murine models (6–8). Chronic corticosterone (CORT) administration, in particular, represents a tractable way to address emotional and cognitive impairments consequence of chronic distress exposure (9). Besides, the CORT model allows examining the direct influence of glucocorticoids on the emergence of a robust pathological phenotype. Corticosterone-treated animals show various signs of disrupted reward-processing or anhedonia-like behaviour, such as reduced preference for palatable solutions or decreased self-directed behaviours like grooming (10–13). Consistent with the scientific literature (13–15), we highlighted high-order cognitive impairment in CORT-treated mice and, most importantly, disruption of optimal decisional processing in a dynamic and uncertain environment (7). More recently, we further demonstrated that various motivational components, i.e. behavioural initiation, effort allocation, and hedonic appreciation and valuation, were altered in CORT-treated mice, and that their behaviour was reminiscent of a typical depressive phenotype (6). Moreover, these animals presented a dampened pattern of neuronal activation (assessed through FosB immunostaining) in the anterior insular cortex (aIC) and the basolateral amygdala (BLA), key telencephalic brain regions involved in appetitive and consummatory motivational processing. To a lesser extent, alterations were also found in the dorsal part of the bed nucleus of the stria terminalis (dBNST) and in the ventral tegmental area (VTA), brain regions known to be involved in the modulation of motivational processes (16,17). However, chronic neural activation in all hese regions failed at predicting the animals’ behavioural output while evaluated in a motivational operant paradigm (6).

To date, scientific evidence strongly indicates that the incorporation of biological factors other than neural activity is necessary in order to understand the biological mechanisms at the origin of stress-related disorders such as major depression and anxiety. Recently, the study of the role of the neurovascular unit (NVU), a concept representing the structural and functional multicellular relationship between the neural tissue and the blood (18), has emerged as a promising approach. Hence, through the study of the NVU components [named neurons, glial cells (perivascular astrocytes and microglia), endothelial cells, pericytes and ependymal cells] and of their intimate associations, the role of this single functioning unit on the emergence of emotional and cognitive dysfunction is being investigated. Although it is still unclear whether neurovascular dysfunction is a cause or a consequence of a particular pathological condition, it has been highlighted in several neurological and neuropsychiatric afflictions (19,20).

Alterations at the NVU level are varied, affecting microvessel architecture, tight junctions (TJ) and the transport function of brain barriers, the blood-brain barrier (BBB) in particular (21,22). Microglial ans astrocytic activity may also contribute to the propagation of neuroinflammatory markers that would alter BBB integrity. In the particular case of anxio-depressive disorders, pro-inflammatory cytokines such as tumour necrosis factor α (TNFα) and interleukin 1β (IL-1β), have been shown to increase BBB permeability in a dose-dependent manner by inducing the expression of intercellular adhesion molecule 1 (ICAM-1) on the luminal surface of BBB endothelial cells in humans and animal models (23,24).

As stated by the increasing scientific literature about the subject, we hypothesize that the development of anxio-depressive behaviour and depressive-related loss of motivation are grounded on modifications of the NVU including BBB integrity. More precisely, we hypothesize that anatomo-functional alterations of glial cells and microvessels, and particularly of TJ, will explain behavioural motivational deficits in mice chronically exposed to exogenous glucocorticoids administration.

The present study aims to determine the role of the NVU in the emergence of motivational deficits consequence of chronic distress. For that, we used available brain samples from mice chronically treated with CORT whose motivation and neuronal activity had been previously evaluated (first data published at (6)), in order to evaluate NVU modifications through immunofluorescence approaches. We also used a new cohort of mice exposed to the same stress protocol to evaluate their BBB permeability. A graphical summary of the experimental design is presented in **Figure 1**. The final goal of our study is to shed light into the relationship between neurovascular alterations and motivational dysfunction relevant in the neurobiological context of neuropsychiatric disorders.

**Figure 1.**
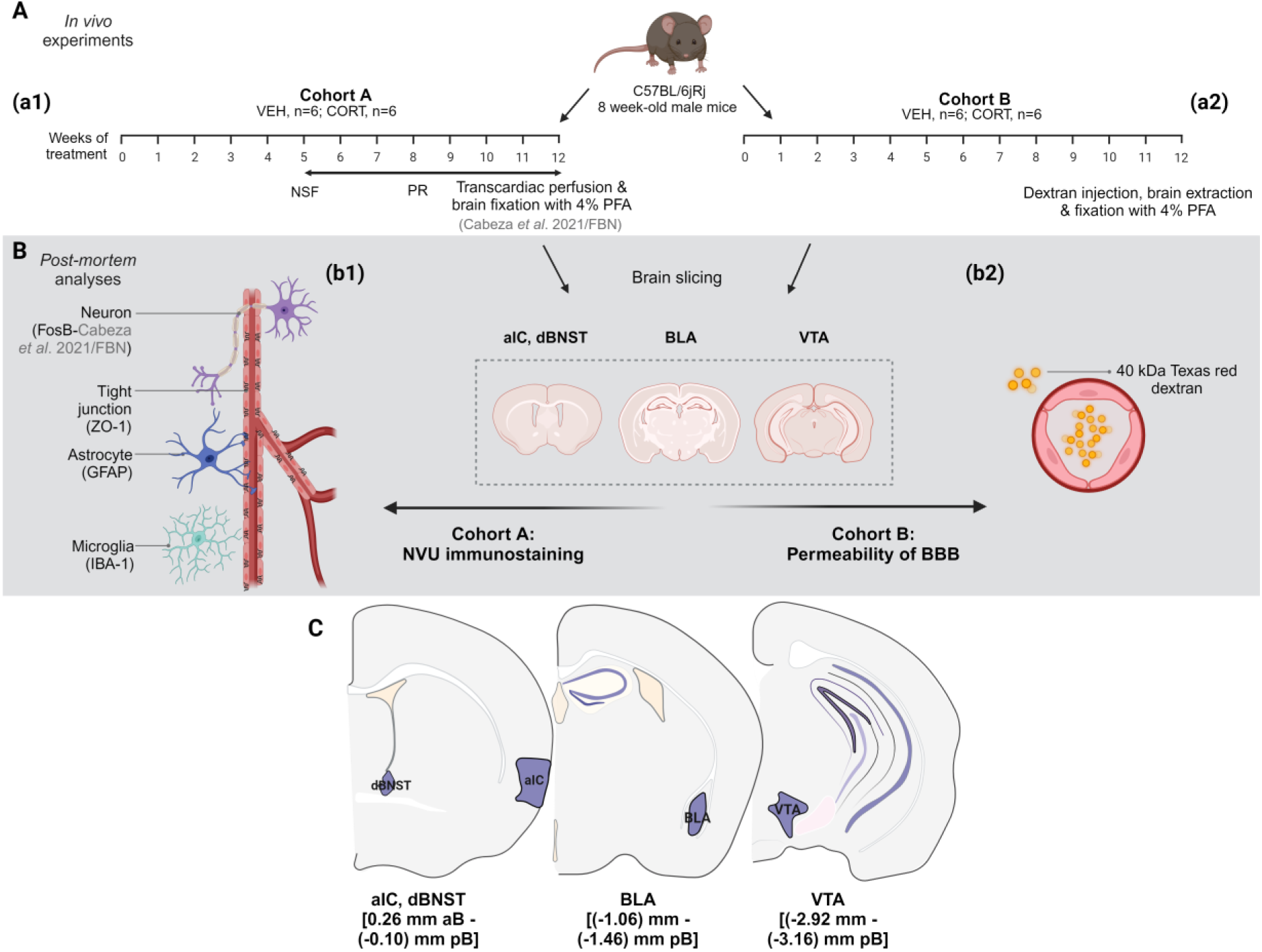
Experimental design. **(A)** Data from two cohorts of adult C57BL/6jRj male mice were used in order to evaluate the role of the neurovascular unit (NVU) in chronic distress-related loss of motivation. **(a1)** Animals from cohort A were differentially treated with either vehicle (VEH) or a corticosterone (CORT) solution for 5 weeks before their motivational state was tested through two behavioral paradigms: the novelty suppressed feeding (NSF) task and an operant progressive ratio (PR) schedule of reinforcement task. After that, animals were euthanized, transcardially perfused with a 4% paraformaldehyde (PFA) solution and their brains collected and processed for immunostaining. **(a2)** Animals from cohort B were differentially treated with VEH or CORT solutions during 12 weeks before being perfused intravenously with fluorescent Texas Red 40 kDa Dextran. Animals were euthanasized immediately after by cervical dislocation under isoflurane anesthesia, and their brains were collected, fixed by immersion in 4% PFA and processed for fluorescence microscopy. **(B)** Four different brain regions involved directly or indirectly in motivational processing (i.e. anterior insular cortex –aIC, dorsal part of the bed nucleus of the stria terminalis –dBNST, basolateral amygdala –BLA, and ventral tegmental area –VTA) were selected in order to study **(b1)** NVU integrity and **(b2)** blood-brain barrier (BBB) permeability in the two experimental conditions. **(b1)** Immunostaining targeting tight junction vessel specific ZO-1, astrocytic specific GFAP and microglial specific IBA-1 were used to evaluate NVU integrity. **(b2)** Levels of Dextran extravasation served as indicators of changes in BBB permeability. **(C)** Anatomical targets and Bregma coordinates of the brain sections selected for immunofluorescence staining of neurovascular components and blood-brain barrier permeability evaluation.

## Methods

Our study of the NVU involvement in the emergence of motivational deficits consequence of chronic distress was attained through (i) a new *post-mortem* immunofluorescence approach and (ii) an *in vivo* line of action.

### Animals

A total number of 24 (Cohort A: n=12; Cohort B: n=12) 6-to 8-week old male C57BL/6jRj mice (*Ets Janvier Labs*, Saint-Berthevin, France) were used in this study. Animals were group-housed and maintained under standard environmental conditions (12 h light/dark cycle; temperature: 22±2 °C; humidity: 55±10%) and had *ad libitum* access to bottles containing vehicle (VEH) or treatment (CORT) (see **Pharmacological treatment** section).

Available brain samples belonged to a cohort of 12 animals (Cohort A: VEH, n=6; CORT, n=6) that had previously been behaviourally evaluated from the sixth week of differential treatment. They confirmed the emergence of motivational deficits while tested in an operant progressive ratio (PR) schedule of reinforcement task evaluating effort allocation (see **Additional information** for more details). As described by Cabeza and colleagues (6), animals were sacrificed after the behavioural testing and brains processed for immunostaining (see **Animals sacrifice and brain sampling** section).

A second cohort of 12 animals was pharmacologically treated for 12 weeks (Cohort B: VEH, n=6; CORT, n=6) before being perfused intravenously with fluorescent Texas Red 40 kDa Dextran (3 mg/kg; D1829, *Fisher Scientific*) as described in previous studies (25,26) and aiming at studying the BBB permeability (see **Animals sacrifice and brain sampling** section).

### Pharmacological treatment

Chronic distress in mice was experimentally induced by sustained oral CORT administration (CORT,-4- Pregnene-11b-diol-3,20-dione-21-dione, *Sigma-Aldrich*, France). Corticosterone was dissolved in a vehicular solution (VEH, 0.45% hydroxypropyl-β-cyclodextrin- βCD, *Roquette GmbH*, France) and administered through the drinking water (35 µg/mL equivalent to 5mg/kg/day). The CORT treatment, as well as the VEH solution, were freshly prepared twice a week as previously described (7,11).

### Animals sacrifice and brain sampling

After an intraperitoneal (IP) injection of pentobarbital (55 mg/kg, Exagon®, *Med’Vet*, France), animals from the Cohort A were transcardially perfused with first 0.9% NaCl and then with 4% ice-cold paraformaldehyde solution (PFA, *Roth®*, Karlsruhe, Germany; in 0.1 M phosphate buffer -PB, pH 7.4), and subsequently postfixed overnight in the same fixative at 4°C. Finally, brains were cryoprotected by immersion in a 15% sucrose solution (D(+)-Saccharose, *Roth®*, Karlsruhe, Germany; in 0.1 M PB) for 24 h at 4 °C, and frozen by immersion in isopentane (2-methylbutane, *Roth®*, Karlsruhe, Germany). Before processing, brains were sliced in 30-µm-thick coronal sections.

Animals from Cohort B were euthanized by cervical dislocation under isoflurane anaesthesia, 20 min after Dextran perfusion. They were then rapidly decapitated, and the brains were extracted and fixed in 4% PFA by immersion during 24 h at 4°C. Next, brains were cryoprotected by immersion in a 15% sucrose solution overnight and frozen by immersion in isopentane. Immediately before observation, 14-µm-thick coronal brain sections were obtained and directly mounted on gelatine-coated slides using a cryostat.

### Immunofluorescence staining and imagery

Brain samples of the Cohort A were immunostained in order to evaluate the influence of chronic distress on the NVU. According to the neural activity modifications previously observed in this cohort of animals (see (6)), and to their role on motivated behaviors and chronic distress, four brain regions were selected for this purpose: the anterior insular cortex (aIC) and the dorsal region of the bed nucleus of the stria terminalis (dBNST) at coordinates (0.26) mm anterior to Bregma (aB) – (0.10) mm posterior to Bregma (pB); the basolateral amygdala (BLA), at (1.06–1.46) mm pB; and the ventral tegmental area (VTA) at (2.92-3.16) mm pB, according to Franklin and Praxinos (27) (see **Figure 1C**). Multiple other brain regions were secondarily analysed and the results are available in **Additional information, Figures 1-2**.

Once mounted on gelatine-coated slides, the sections followed an antigen retrieval protocol aiming at maximizing antigenic site exposure for antibody binding. Slides containing the sections were immersed for 40 min in 10 mM citrate buffer pH 6 (sodium citrate, *Sigma Aldrich*, Germany) at 96°C in a water bath. The buffer was then left to cool down at room temperature before removing the samples. After washing, the sections were incubated with the primary antibodies as follows: co-staining IBA-1 – ZO-1 incubation during 24 h at room temperature (1:1000, ab178846, rabbit anti-IBA-1, *Abcam;* 1:200, sc-33725, rat anti-ZO-1, *Santa Cruz Biotechnology*); and GFAP incubation during 24 h at room temperature (1:1000, AB5804, rabbit anti-GFAP, *EMD Millipore*). Next, the sections were incubated with the secondary antibodies overnight at room temperature (1:500; Alexa Fluor^TM^ 488, donkey anti-rabbit IgG (H+L); Cy3^TM^, goat anti-rat IgG (H+L)). Finally, sections were washed and coverslipped with mounting media (40% PB, 60% glycerol, *Roth®*, Karlsruhe, Germany; DAPI Fluoromount-G®, *SouthernBiotech*, Birmingham, USA). Photomicrographs of brain structures for analysis were acquired using 10x or 20x objectives of an Olympus microscope Bx51 equipped with an Olympus DP50 camera. *ImageJ* Fiji software (National Institute of Health, Bethesda, Maryland, USA, http://imagej.net/ij/ (28)) was used to quantify the antigen-related fluorescence in terms of intensity and occupied surface, and in the particular case of IBA-1, also to count the number of cells per unity of surface. For final sample sizes, see **Additional Table 1**.

Brain samples from Cohort B, after being mounted on gelatine-coated slides and coverslipped with mounting media, were observed and pictures of the regions of interest were taken with a ZEISS Axio Imager.Z2 microscope, equipped with ApoTome.2 and a Camera ORCA-Flash4.OLT (*Zeiss*, Germany). Photomicrographs were acquired using 10x and 40x objectives, and processed to get maximal orthogonal intensity projections, and to reproduce the neural environment in a three-dimensional construct. The *Zen2 pro* (blue edition) version 2.0.0.0 software (©*Carl Zeiss Microscopy GmbH*, 2011) was used to quantify vascular Dextran diffusion through the measure of the ratio between intra- and extravascular fluorescence intensity.

### Data and statistical analyses

The results are presented as means ± SD. The statistical analyses were conducted using STATISTICA 10 (*Statsoft*, Palo Alto, United States) and figures were designed using GraphPad Prism version 10.2.1 for Windows (*GraphPad Software*, Boston, Massachusetts, USA, www.graphpad.com). The nature of the data sets was verified using Shapiro-Wilk and Levene’s tests to respectively study the data sets’ normality of distribution and the homogeneity of variance. Since data did not meet assumptions for parametric analysis, data from the two experimental conditions were compared using Mann-Whitney U (MWU) tests. The relationship between the behavioural scores and the selected anatomical markers (FosB, IBA-1, ZO-1 and GFAP expression) in the brain regions of interest was studied using principal component analysis (PCA), which allow maximizing the explanatory variance across the measured parameters. Dimensional relationships between behavioural (i.e. motivational) and neurobiological scores (NVU and FosB parameters) were analysed using Pearson correlations. For all analyses, the significance level was set as p<0.05 and dependencies with p≤0.1 are interpreted as trends.

## Results

### Chronic distress alters glial components of the NVU in the anterior insular cortex (aIC) and in the bed nucleus of the stria terminalis (dBNST)

The impact of chronic exposure to exogenous glucocorticoids on three different NVU components, i.e. TJ, microglia and astrocytes, was studied through, respectively, the expression of the specific markers ZO-1, IBA-1 and GFAP in the brain regions of interest.

#### Results from the aIC

Concerning the relative expression of these NVU components, here called relative fluorescence intensity, no effect was observed in the aIC as illustrated in **Figure 2A** (mean fluorescence intensity of ZO-1 ± SD (%), VEH: 100.00±17.17; CORT: 88.54±10.42; IBA1, VEH: 100.00±10.04; CORT: 101.90±6.30; GFAP, VEH: 100.00±17.85; CORT: 78.61±26.58) [VEH vs CORT, MWU; ZO-1: Z=1.6, p=0.11; IBA-1: Z=-0.5, p=0.58; GFAP: Z=1.4, p=0.15]. However, in terms of the surface occupied by these markers, a significant increase was observed for GFAP in the aIC (mean of occupied surface ± SD (%); VEH: 2.94±0.60; CORT: 5.23±2.06) [Z=-2.2, p=0.025], but not for ZO-1 (VEH: 19.25±7.20; CORT: 23.78±6.75) [Z=-1.4, p=0.15], neither for IBA-1 (VEH: 8.00±1.89; CORT: 6.44±1.09) [Z=1.3, p=0.20] (**Figure 2B**).

**Figure 2.**
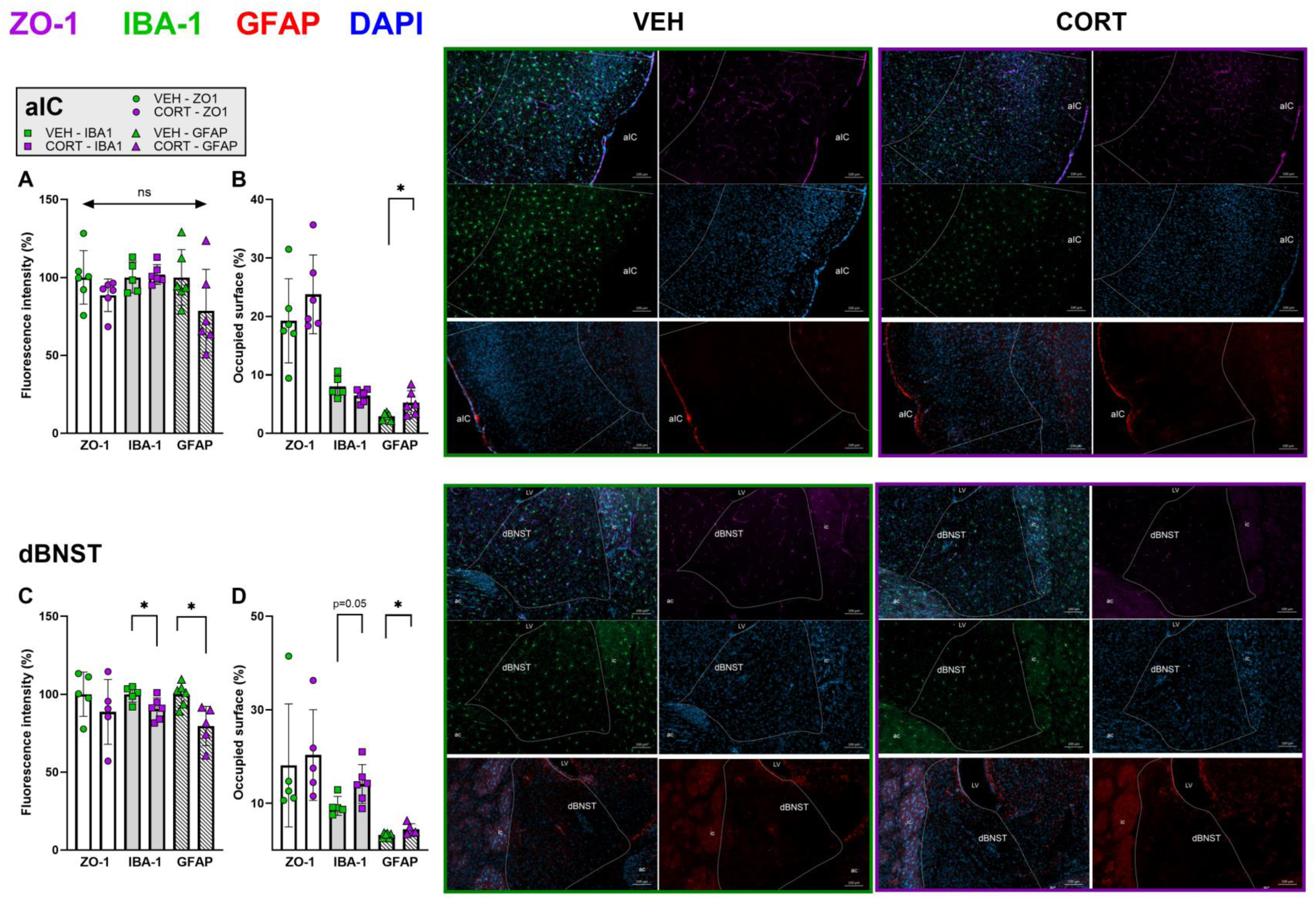
Chronic CORT administration impacts glial components of the NVU in the anterior insular cortex and in the bed nucleus of the stria terminalis. The effect of chronic distress on the NVU integrity was studied through the expression of specifics biomarkers of three different components: ZO-1 (violet) as vessel specific tight junction element; IBA-1 (green) for microglial cells; and GFAP (red) for astrocytes. In the anterior insular cortex (aIC), **(A)** no effect of the CORT treatment was observed in what concerns the relative fluorescence intensity (i.e. relative expression) of the NVU components. However, **(B)** the surface occupied by GFAP in the aIC was significantly increased in CORT-treated animals compared to their control counterparts. No other effects were observed, neither for ZO-1 nor for IBA-1. In the dorsal region of the bed nucleus of the stria terminalis (dBNST), **(C)** IBA-1 and GFAP relative expressions were significantly decreased in mice treated with CORT, **(D)** while their respective occupied surfaces were significantly increased. No effect of the CORT treatment on ZO-1 was observed. Illustrative photomicrographs of ZO-1, IBA-1 and GFAP immunostaining for the experimental conditions VEH vs. CORT, taken with a ZEISS Axio Imager Z2 microscope, equipped with ApoTome.2 and a Camera ORCA-Flash4.OLT (*Zeiss*, Germany). Values are mean ± SD (n=5-6 per group). *, p<0.05; ns, not significant.

#### Results from the dBNST

The glial NVU components were significantly impacted in the dBNST, while ZO-1-related measures were similar between the experimental conditions (**Figures 2C-D**). More specifically, IBA-1 and GFAP relative expressions were decreased in CORT animals (IBA-1, VEH: 100.00±2.29; CORT: 90.61±2.89; GFAP, VEH: 100.00±3.03; CORT: 79.64±6.62) [IBA-1: Z=2.2, p=0.028; GFAP: Z=2.4, p=0.018], and their respective occupied surfaces increased (IBA-1, VEH: 9.46±2.03; CORT: 14.14±4.15; GFAP, VEH: 3.23±0.50; CORT: 4.48±1.16) [IBA-1: Z=-2.0, p=0.045; GFAP: Z=-2.4, p=0.018]. No effect of the CORT treatment was observed in what concerns to ZO-1 in the dBNST (fluorescence intensity, VEH: 100.00±6.35; CORT: 88.69±9.29; occupied surface, VEH: 18.12±13.16; CORT: 20.34±9.70) [fluorescence intensity: Z=0.9, p=0.35; occupied surface: Z=-0.9, p=0.35].

### Altered relative expression and surface coverage of vessel-specific ZO-1 in the basolateral amygdala (BLA) after chronic distress

#### Tight junctions’ alterations

Chronic CORT administration significantly impacted the vessel TJ component ZO-1 in the BLA as illustrated in **Figure 3**. CORT-treated animals present significantly decreased values of ZO-1 fluorescence intensity (85.07±7.95) in the BLA compared to VEH-treated animals (100.00±12.05) [MWU: Z=2.2, p=0.025], while the surface occupied by it is significantly increased (VEH: 12.56±2.68; CORT: 17.48±2.76) [Z=-2.4, p=0.016]. These bidirectional results suggest a reorganization of TJ-specific ZO-1 in animals under chronic distress (see complementary results in **Additional Figure 3**). Besides, irregularities and interruptions can be observed in the vessels’ ZO-1 staining of CORT-treated individuals, and which are not detected in control animals (**Figures 3C2-D2**).

**Figure 3.**
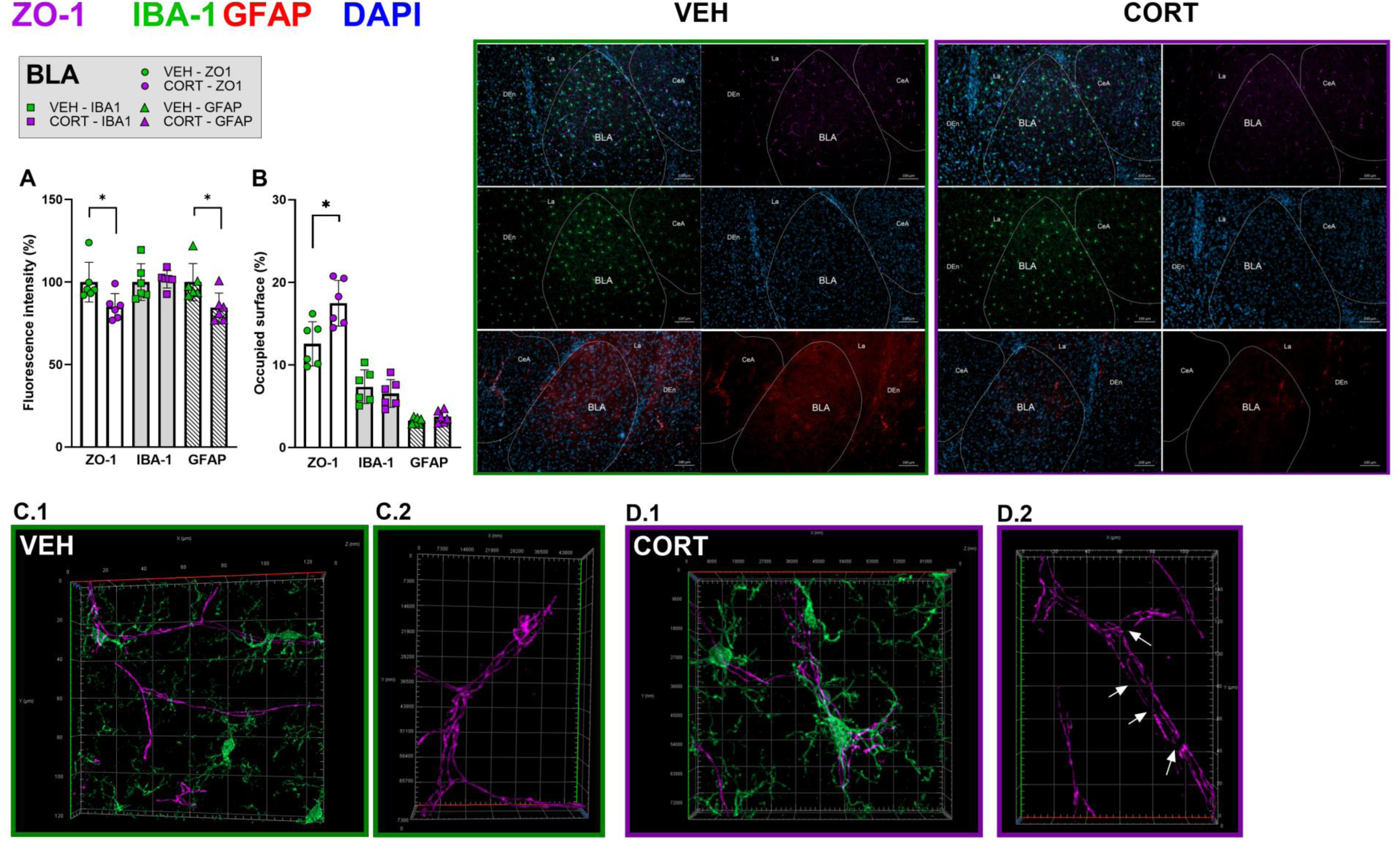
Chronic CORT administration alters relative expression and surface coverage of vessel-specific ZO-1 in the basolateral amygdala. **(A)** The relative fluorescence intensity (i.e. relative expression) of ZO-1 and GFAP in the basolateral amygdala (BLA) were significantly decreased in CORT-treated mice compared to VEH-treated animals. **(B)** However, chronic CORT administration only impacted the surface occupied of ZO-1, which was significantly increased in distressed mice. The values obtained for surface occupied by GFAP in the BLA were similar in VEH- and CORT-treated animals. No effect of the CORT- treatment was observed on IBA-1-related measures in the BLA. Illustrative photomicrographs of ZO-1 (violet), IBA-1 (green) and GFAP (red) immunostaining for the experimental conditions VEH vs. CORT, taken with a ZEISS Axio Imager Z2 microscope, equipped with ApoTome.2 and a Camera ORCA-Flash4.OLT (*Zeiss*, Germany). Illustrative 3D reconstructions of BLA microglial cells and vessels of control **(C.1-2)** and CORT-treated **(D1-2)** animals. Arrows in **D.2** point to observed irregularities in ZO-1 related microvessel scaffolding. Values are mean ± SD (n=5-6 per group). *, p<0.05.

#### Astrocytes and microglia adjustments

A significant decrease in GFAP relative expression was observed in the BLA of CORT-treated animals (84.54±8.87) compared to their control counterparts (100.00±11.26) [Z=2.1, p=0.037], but the occupied surface by this astrocytic component was similar between the experimental conditions (VEH: 3.26±0.39; CORT: 3.74±0.71) [Z=-1.3, p=0.20]. No effect of the CORT-treatment was observed on IBA-1 relative expression and occupied surface in the BLA (fluorescence intensity, VEH: 100.00±11.13; CORT: 101.70±5.27; occupied surface, VEH: 7.37±2.04; CORT: 6.55±1.68) [fluorescence intensity: Z=- 0.8, p=0.42; occupied surface: Z=0.80, p=0.42].

### Chronic distress alters vessel-specific ZO-1 relative expression in the ventral tegmental area (VTA)

#### Tight junctions’ alterations

A significant decrease of ZO-1 relative fluorescence intensity in the VTA of CORT-treated animals (75.95±13.93) compared to animals from the control group (100.00±8.78) [MWU: Z=2.4, p=0.014], as well as a tendency to an increase in the surface occupied by it (VEH: 3.94±1.00; CORT: 6.85±3.12) [Z=- 1.6, p=0.10] (**Figure 4A-B**), similar to BLA observations.

**Figure 4.**
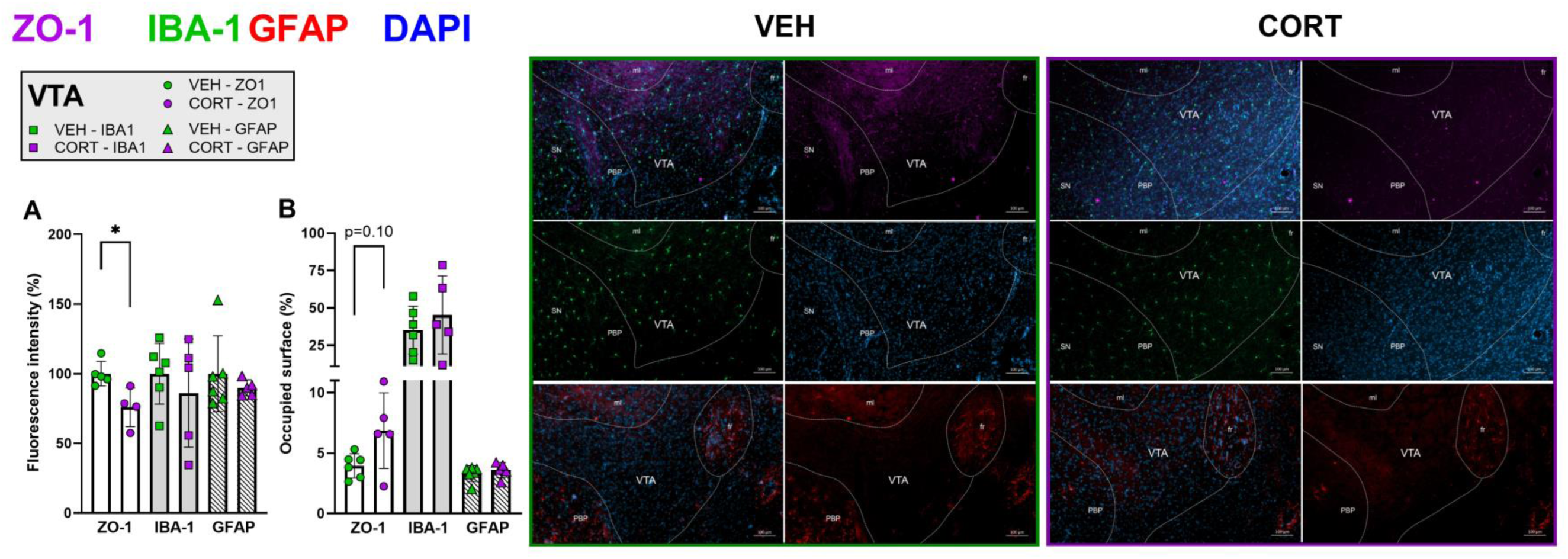
Chronic distress alters vessel-specific ZO-1 relative expression in the ventral tegmental area. **(A)** The relative fluorescence intensity (i.e. relative expression) of ZO-1 was found significantly decreased in the ventral tegmental area (VTA) of animals chronically treated with CORT compared to their control counterparts. Concerning IBA-1 and GFAP relative expression, no differences were observed between the experimental groups. **(B)** The VTA surface occupied by ZO-1 was found to tend to an increase in CORT-treated animals compared to control mice, without reaching significance. No other effects of the CORT treatment were observed, neither for IBA-1 nor for GFAP-related measurements. Illustrative photomicrographs of ZO-1 (violet), IBA-1 (green) and GFAP (red) immunostaining for the experimental conditions VEH vs. CORT, taken with a ZEISS Axio Imager Z2 microscope, equipped with ApoTome.2 and a Camera ORCA- Flash4.OLT (*Zeiss*, Germany). Values are mean ± SD (n=5-6 per group). *, p<0.05.

#### Astrocytes and microglia adjustments

No alterations were observed, neither for IBA-1-related measurements (VEH, fluorescence intensity: 100±21.78; occupied surface: 35.11±16.05; CORT, fluorescence intensity: 86.10±38.85: occupied surface: 45.32±26.13) [fluorescence intensity: Z=0.5, p=0.58; occupied surface: Z=-0.7, p=0.47], nor for GFAP-related measurements (VEH, fluorescence intensity: 100.00±27.25; occupied surface: 3.30±0.65; CORT, fluorescence intensity: 89.90±5.73: occupied surface: 3.59±0.63) [fluorescence intensity: Z=0.2, p=0.86; occupied surface: Z=-0.7, p=0.47] within the VTA.

### Chronic CORT administration does not impact microglial density in the brain regions showing altered patterns of neural activation

In order to further evaluate the effect of sustained exogenous glucocorticoids exposure on microglial reactivity, the local density of this cellular population was compared between experimental conditions. Chronic CORT administration did not significantly affect microglial density in the aIC (mean number of IBA-1 positive cells ± SD, VEH: 262.29±58.02; CORT: 241.84±34.14) [MWU: Z=0.5, p=0.58], the dBNST (VEH: 166.07±11.98; CORT: 157.74±22.35) [Z=0.9, p=0.39] or the BLA (VEH: 281.75±44.33; CORT: 236.11±38.70) [Z=1.5, p=0.14], and only a tendency to a decreased density was observed in the VTA of CORT-treated animals (144.00±19.06) compared to their control counterparts (188.80±43.32) [Z=1.7, p=0.082] (**Additional Figure 4A**). However, a general pattern of decreased microglial density was observed through the evaluation of other brain regions, as illustrated in **Additional Figure 4B**.

### Chronic CORT administration alters BBB permeability in the anterior insular cortex, the basolateral amygdala and the ventral tegmental area

Aiming at evaluating whether and to what extent the permeability of the BBB is impacted by sustained CORT administration and by the alterations induced on ZO-1 and GFAP expression and/or distribution, we used Texas Red 40 kDa Dextran as an exogenously administered fluorescent marker of vascular integrity. Vascular areas outlined by Dextran were subsequently compared between the experimental conditions in the four brain regions of interest (i.e. aIC, dBNST, BLA and VTA), revealing significant differences in the ratio between the detected fluorescence outside vs. inside the vessels in the aIC (mean ratio of out/in fluorescence intensity ± SD, VEH: 0.57±0.06; CORT: 0.68±0.06) [MWU: Z=-2.3, p=0.020], the BLA (VEH: 0.66±0.08; CORT: 0.76±0.07) [Z=-2.1, p=0.037] and the VTA (VEH: 0.72±0.10; CORT: 0.86±0.02) [Z=-2.2, p=0.030]. No difference in the outside/inside ratio was observed in what concerns the dBNST (VEH: 0.65±0.07; CORT: 0.68±0.07) [Z=-1.0, p=0.33]. Hence, the higher out/in Dextran-related fluorescence ratios characterizing CORT-treated individuals indicate increased extravasation of the agent into the brain parenchyma (see **Figure 5**). Interestingly, BBB permeability is only altered in the brain areas where ZO-1 seems delocalized, i.e. in regions where ZO-1 expression inversely correlated with the surface occupied by it (**Additional Figure 3**) [aIC: R^2^=0.84, p<0.0001; BLA: R^2^=0.46, p=0.011; VTA: R^2^=0.94, p<0.0001].

**Figure 5.**
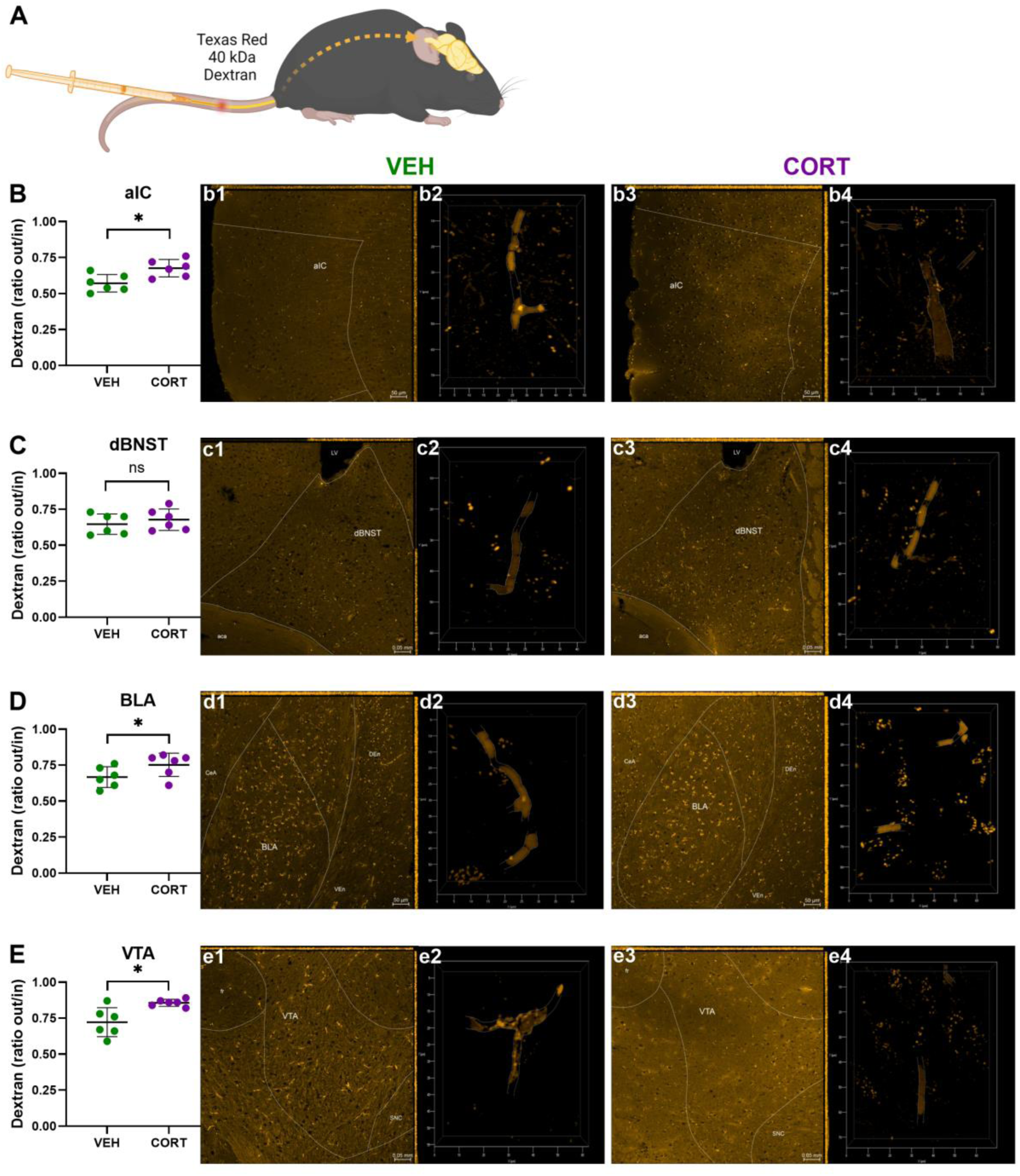
Chronic CORT administration alters neurovascular unit’s permeability in the anterior insular cortex, the basolateral amygdala and the ventral tegmental area. **(A)** Mice from Cohort B were perfused with fluorescent Texas Red 40 kDa Dextran and euthanized after 20 min. Fluorescence intensities directly outside and inside microvessels were calculated from coronal brain sections 14-µm- thick containing the four regions of interest (i.e. anterior insular cortex –aIC, dorsal part of the bed nucleus of the stria treminalis –dBNST, basolateral amygdala –BLA, and ventral tegmental area –VTA). Dextran extravasation was found significantly increased in **(B)** the aIC, **(D)** the BLA and **(E)** the VTA of CORT-treated animals compared to control (VEH) mice. **(C)** No difference between the experimental conditions was observed in the dBNST. **(b1,3; c1,3; d1,3; e1,3)** Illustrative photomicrographs of Dextran-related fluorescence for the experimental conditions VEH vs. CORT in the 4 regions of interest, taken with a ZEISS Axio Imager Z2 microscope, equipped with ApoTome.2 and a Camera ORCA- Flash4.OLT (*Zeiss*, Germany). **(b2,4; c2,4; d2,4; e2,4)** Three-dimensional reconstructions of individual microvessels illustrating differences of Dextran extravasation between experimental conditions. Values are mean ± SD (n=6 per group). *, p<0.05; ns, not significant.

### Neurovascular unit modifications in the ventral tegmental area predict motivation-related behavioural output in distressed mice

In order to further evaluate the putative causal relationship between NVU modifications and motivational deficits subsequent to chronic CORT administration in cohort A, PCA analyses including all parameters under study (relative fluorescence intensity and occupied surface of ZO-1, IBA-1 and GFAP; microglial density; motivation behavioural score “breakpoint” -BP) in the four regions of interest (aIC, dBNST, BLA and VTA) were performed.

Based on the PCA results, four principal components (PC1-4) entirely explain the variance of the data set’s metrics, with the three first PCs (PC1-3) edifying for more than 75% (i.e. 87.13%), as illustrated in **Figure 6A**. The loading values (i.e. correlation values from data vs. eigenvectors) of PC1-3 are plotted as heat-map aiming at graphically unfold the contribution of single variables within single PCs (**Figure 6B**). The yielded 4-factor overall solution obtained in the current study therefore alludes to the close relationship hold by the investigated parameters. With the focus on the motivational data (i.e. variable BP), VTA NVU components strongly correlate with PC1, with PC1 increasing as BP (loading value: - 0.952), ZO-1 fluorescence intensity (–FI: −0.972), microglial density (–MD: −0.964), GFAP FI (−0.863) and IBA-1 FI (−0.845) decrease, and as ZO-1 occupied surface (–OS: 0.924) and GFAP OS (0.841) increase. Furthermore, strong correlations are also evidenced for BLA ZO-1 FI and FosB expression, and for dBNST IBA-1 FI, with PC1 increasing as these parameters decrease (−0.932, −0.818 and −0.884, respectively). Therefore, motivational performances are mainly explained by PC1 and robustly predicted by VTA NVU components. This association is conspicuous in the loading plot (**Figure 6C**), which graphically illustrates the variables’ relationships within PC1 and PC2. As previously indicated in **Figure 6B**, PC2 mostly correlates with BLA MD and IBA-1 FI (0.927 and 0.786, respectively), dBNST FosB expression (−0.926) and aIC ZO-1 FI (0.776). However, the contribution of the variables to PC2 have a poor influence on BP (−0.260) (**Figure 6C**). Complementary linear correlation analyses between motivational scores (BP) and the identified variables most strongly correlating with PC1 reveal an apparent clustering of the data set in VEH- vs CORT-treated individuals (**Figure 6D-I**). Thus, relative expression of ZO-1 in the BLA and VTA, and of IBA-1 in the dBNST, which are decreased in CORT-treated animals, predict motivational performance in the PR paradigm (**Figure 6D,I**). Additional correlations between NVU components and behavioural scores can be found in **Additional Figure 5**.

**Figure 6.**
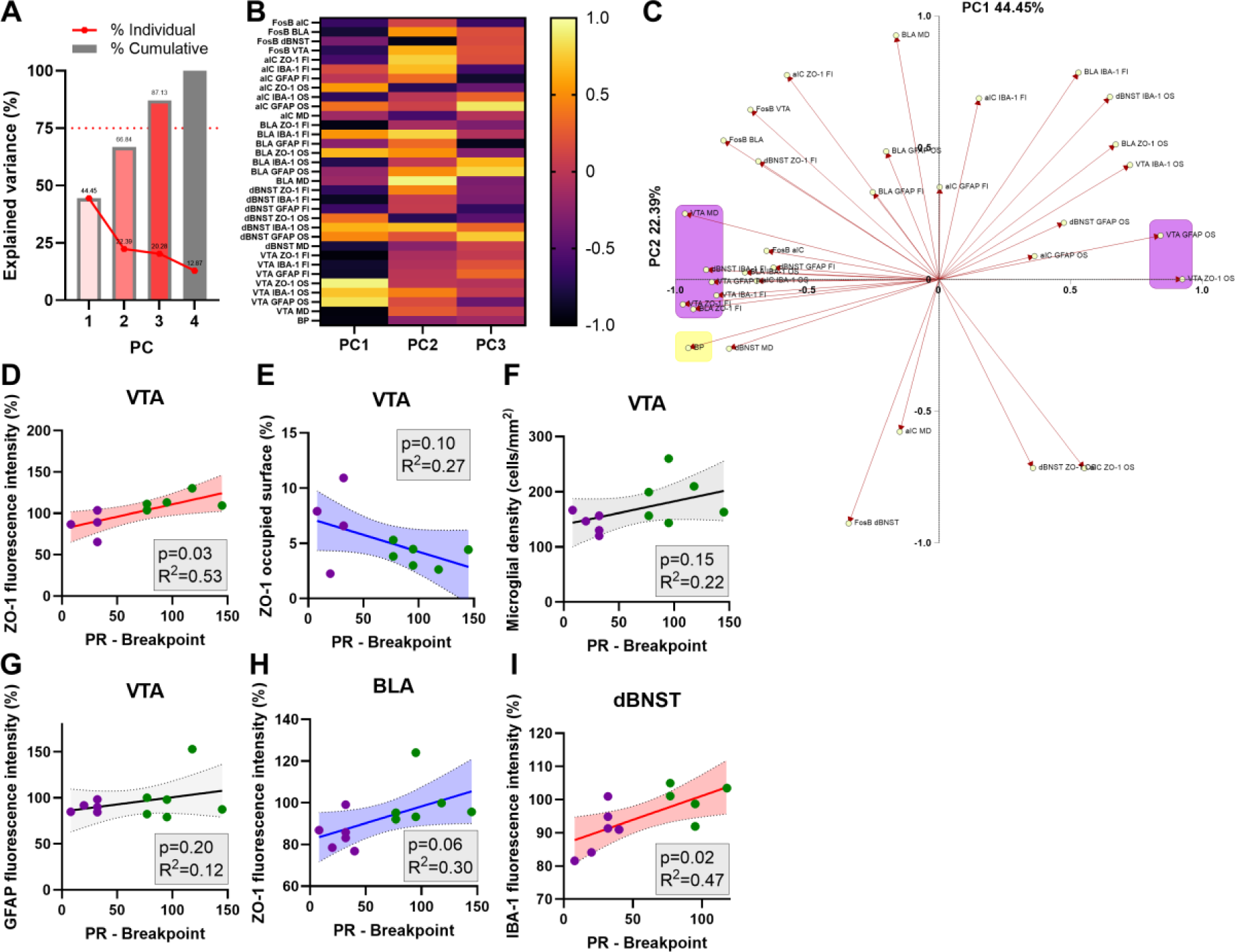
Neurovascular unit modifications in the ventral tegmental area predict motivational performances in distressed mice. Motivational performances from cohort A (breakpoint –BP, i.e. last ratio attained in the progressive ratio –PR– schedule of reinforcement task) were confronted to the neurobiological measurements (relative fluorescence intensity –FI, and occupied surface –OS, of ZO- 1, IBA-1 and GFAP; and microglial density –MD) through a principal component analysis (PCA). **(A)** Four principal components (PC1-4) explain the total variance of the data set, with the three first PCs (PC1- 3) explaining 87.13%. **(B)** The heat-map of the loading values of PC1-3 unfolds a strong contribution of VTA NVU components in the behavioural performance of mice in the motivational task (direct from ZO-1 FI, MD, GFAP FI and IBA-1 FI; indirect from ZO-1 and GFAP OS. Another direct strong contribution was revealed for BLA ZO-1 FI and FosB expression, and for dBNST IBA-1 FI. **(C)** The loading plot illustrates the variables’ relationships within PC1 and PC2, with PC2 mostly correlating with BLA MD and IBA-1 FI, dBNST FosB expression and aIC ZO-1 FI, but poorly influencing motivational performance. **(D-I)** Linear correlations between motivational scores (BP) and variables strongly correlating with PC1 reveal an apparent clustering of the data set in VEH- vs CORT-treated individuals.

## Discussion

The results of the present experimental investigation highlight a relationship between NVU adjustments to chronic distress and poor behavioural performance in mice. Most particularly, observed variations related to TJ-specific ZO-1, which is a key element of the cerebral vessel scaffolding, are associated to an increased BBB permeability in distressed animals and robustly explain motivational deficits assessed through an effort-based operant behavioural paradigm. Thus, to the best of our knowledge, our results show for the first time that emotional-cognitive alterations in mice reminiscent to human neuropsychiatric symptomatology, and in particular to loss of motivation, are largely grounded on neurovascular adjustments within brain centres for motivational processing. Hence, we support that neuronal activity cannot be further taken as an isolated major actor sustaining behavioural performance neither in physiological nor in pathological contexts.

In our evaluation of the impact of chronic distress on mice NVU components, we identified several alterations in the pattern of expression of vessel and glial markers. A major observation concerns the expression of the specific TJ element ZO-1, which was found significantly decreased in the BLA and VTA of CORT-treated animals but not in the aIC. These three brain areas presented increased ratios of Dextran extravasation in distressed animals, which confirms a direct impact of the stress protocol on local BBB permeability. Interestingly, through our immunofluorescence staining analysis, we found that the surface occupied by ZO-1 in the aIC, BLA and VTA negatively correlates with its expression (**Additional Figure 3**). We propose here that chronic distress might induce ZO-1 delocalization from endothelial TJ, contributing this way to an increase in the BBB permeability. In other words, ZO-1 delocalisation, attested by the negative correlation between ZO-1 expression and occupied surface, might better predict BBB integrity/permeability than ZO-1 expression alone in endothelial cells. Delocalization mechanisms of ZO-1 have been previously described, for example, following a calcium switch (i.e. low calcium levels in the cellular medium disrupt cell/cell junctions) in kidney epithelial cells, forcing ZO-1 into the cytoplasm (29). It would be therefore insightful to disclose whether other structural elements of endothelial TJ are also impacted by chronic CORT administration, and might contribute to the altered mice BBB permeability. Along with this idea, altered expression of adhesion molecules in BBB endothelial cells has been proposed to be at the origin of BBB hyperpermeability in depression (30,31). A down-regulation of TJ protein Claudin-5 has been observed in the nucleus accumbens (NAc) and in the medial prefrontal cortex (mPFC) of mice exposed to social defeat stress (2,32). Furthermore, the work of Wu and colleagues (2022) demonstrated a link between Claudin-5 down-regulation and the emergence of anxious-like behaviour in female mice, as well as a decrease of these behaviours in mice under social isolation by overexpressing Claudin-5 in the amygdala (33). Similarly, in the chronic social defeat murine model of depression, the expression of endothelial Claudin-5 is reduced along with abnormal blood vessel morphology in the NAc of stress-susceptible but not resilient mice (32). Altogether, this scientific evidence strongly implies a key role of the integrity of the BBB, and in particular of the endothelial-related vessel scaffolding allowing intimate cellular junctions, in the emergence of emotional-cognitive deficits in animals exposed to stress.

In this study, we also identified distress-induced NVU alterations at the glial level. The expression of the specific astrocytic marker GFAP was significantly decreased in the BLA and the dBNST of CORT-treated mice, while the surface occupied by it was increased in the aIC and the dBNST. After an acute stress, astrocytes are known to regulate BBB permeability via the release of cytokines, including interleukin 1-beta (IL-1β) (34). Besides, in response to IL-1β stimulation, astrocytes release vascular endothelial growth factor (VEGF), a key modulator of NVU function that increase BBB permeability (34). Accordingly, our results suggest that distress impacts astrocytic activity. Interestingly, GFAP expression significantly correlates with the surface occupied by ZO-1 in the BLA (**Additional Figure 6A**), suggesting that it might be indirectly involved in the increase of the BBB permeability in this brain area.

To better understand the relationship between the NVU and the motivational state of control and CORT-treated mice, we used a PCA approach. Thus, we tested whether NVU components, including neuronal activation, influence the motivational profiles of VEH and CORT-treated animals. Despite the small sample size available, the PCA results yielded a 4-factor full solution for the metrics data set. This particular fact (i.e. 100% of the variance of the data set is explained by 4 PCs) highlights the close relationship and the substantial contribution of each of the parameters evaluated to the study. Notably, our PC1, which is characterized by prominent loadings for VTA NVU components, outstandingly predicts animals’ behavioural performance in the PR paradigm. In particular, ZO-1 adjustments in terms of expression and distribution in the VTA and the BLA following distress may be key determinants at the origin of the motivational deficits, rather than neural activity. Besides, the neurovascular adjustments observed in the aIC and the dBNST seem to play a secondary role in the behaviour evaluated. Noteworthy, the motivational behaviour here evaluated involved rewarding food intake. Reward and food intake information processing relay on several brain areas and circuits, among them the VTA, BLA, dBNST and aIC. This complexity may explain why neuronal activation of single regions failed to translate the motivational performance observed through de use of lineal correlation models (6). However, our PCA analysis, which took into consideration FOSB expression levels from the 4 brain areas, confirms that neural activity alone does not explain the motivational behaviour of VEH and CORT-treated mice.

Our results clearly evidence, for the first time, a strong relationship between the integrity of the TJ of brain capillary endothelial cells and motivational behaviour of control and distressed mice. Although the mechanistic process that leads to this NVU-behaviour relationship deserves to be studied in depth, our results suggest the existence of close communications between NVU cells at the local level and/or between distinct brain areas. It has been recently demonstrated that brain cells, including microglia, astrocytes and endothelial cells can communicate each other by forming tunnelling nanotubes (for review see (35)). To our knowledge, however, no interregional relationships have been identified between components of the NVU. Hence, we studied whether astrocytes, microglia and brain endothelial cells may interact with each other along the four brain regions studied (i.e. aIC, dBNST, BLA and VTA). We found several significant linear correlations between these areas (**Additional Figure 6**). Interestingly, these correlations mainly evidence a microglial-endothelial cells relationship between VTA/BLA, VTA/dBNST, dBNST/BLA and BLA/aIC (see summary in **Additional Figure 6B**). In line with our observations within the BLA, these results suggest an enhanced relationship between microglial cells and brain microvessels in CORT-treated animals (**Figure 3C1-D1**). Noticeable, the regions concerned by the IBA-1/ZO-1 relationship match with those where a neurons/neurons relationship was identified. In this line, optogenetic stimulation approaches have demonstrated that VTA projecting to BLA dopamine neurones are involved in the modulation of anxious-like behaviour in mice (36). Chemogenetic manipulations have evidenced that the VTA-BLA-NAc neural circuit regulates reward valuation and motivated behaviour (37). It is known that the BLA is involved in the encoding of motivationally-relevant representations of specific outputs from instrumental associations (38) and that acts as sensor of incentive values changes (39,40). The aIC-amygdala connectivity seems to be essential for valence processing (41) and for shaping emotional output and subsequent behaviour (42). The dBNST shares a common pathway with the BLA, and participates in the integration of the stress information, as well as in the regulation of food intake (43,44). Taken in a whole, our data suggest relevant interregional communication pathways closely involving neurons and other NVU components, and leading to adaptive motivational behaviour in the CORT murine model of chronic distress.

Our findings expand on our previous work and propose specific biological targets for the causal relationship between chronic distress and emotional-cognitive dysfunction. However, the neuroanatomy of these brain regions is complex (e.g. the VTA is 60% dopaminergic, 35% GABAergic and 5% glutamatergic), several pathways may influence motivated behaviour, and little is known about the interregional effects of local neurovascular adjustments. Future experiments should address these questions.

Finally, it is likely that local neural pattern activation following NVU adjustments represents a dynamic balance between damage and repair mechanisms. Hence, future investigation should target discriminatory biomarkers of pro-dysfunction and pro-repair signalling mechanisms in order to improve precision medicine strategies.

## Conclusions

The NVU is a complex concept representing a variety of cells and interactions that plays a central role on cerebral homeostatic control. Our results confirm that NVU components are highly reactive to stress, and that their integrity account for the quality of an individual behaviour. We present evidence placing the BBB in a privileged position for the modulation of the emotional-cognitive behaviour of individuals under stressful conditions. More specifically, we demonstrated that the integrity of TJ between BBB endothelial cells is determinant in this context. However, and despite recent research on NVU responsiveness to distress, further investigation is needed in order to shed light into the underlying mechanisms at the origin of emotional-cognitive dysfunction.

## List of abbreviations

aIC: Anterior insular cortex
BBB: Blood-brain barrier
BLA: Basolateral amygdala
BP: Breakpoint
CORT: Corticosterone
dBNST: Dorsal region of the bed nucleus of the stria terminalis
FI: Fluorescence intensity
GFAP: Glial fibrillary acidic protein
IBA-1: Ionized calcium-binding adapter molecule 1
ICAM-1: Intercellular adhesion molecule 1
IL-1β: Interleukin 1 beta
IP: Intraperitoneal
MD: Microglial density
mPFC: Medial prefrontal cortex
MWU: Mann-Whitney U
NAc: Nucleus accumbens
NSF: Novelty suppressed feeding
NVU: Neurovascular unit
OS: Occupied surface
pB: Posterior to Bregma
PCA: Principal component analysis
PFA: Paraformaldehyde
PR: Progressive ratio
TJ: Tight junction
TNF-α: Tumour necrosis factor α
VEGF: Vascular endothelial growth factor
VEH: Vehicle
VTA: Ventral tegmental area
ZO-1: Zonula occludens 1
βCD: Hydroxypropyl-β-cyclodextrin

## Declarations

### Ethics approval and consent to participate

All animal procedures were conducted in accordance with the Guide for the Care and Use of Laboratory Animals (NIH), the Animal Research: Reporting of In Vivo Experiments (ARRIVE) guidelines, and the European Union regulations on animal research (Directive 2010/63/EU). They were approved by the University of Franche-Comté Animal Care and Use Committee (CEBEA-58).

### Consent for publication

Not applicable.

### Availability of data and materials

The datasets used and/or analyzed during the current study are available from the corresponding author on reasonable request.

### Authors’ contributions

LC and FB conceived the experiments. LC, DM, BR and CH carried out the experiments. LC, DM, AE and FB analysed data. LC, DM, GBC, AE and FB wrote the paper. YP, EH and the rest of the authors critically revised the work and approved the version to be published.

## Acknowledgements

We thank PhD Pierre-Yves Risold (LINC) for his helpful insight during the immunofluorescence experiments, and PhD Sophie Croizier and her team (University of Lausanne) for their help in the acquisition of microscopic images.

