## Additional information for "Neurovascular unit adjustments following chronic distress explain motivational deficits in mice"


### Additional Information

**Behavioral protocol *as described by Cabeza et al., 2021 (doi: 10.3389/fnbeh.2021.717701).***

***Progressive ratio schedule of reinforcement task (PR).***

Operant behavior was evaluated using five identical two-hole-operant chambers (Med Associates, Hertfordshire, United Kingdom), individually housed inside sound-attenuating cubicles with ventilating fans. The chambers were connected to a PC computer with MedPC-5 software via a Smart Control interface cabinet and programs were written in Medstate notation code.

A habituation period and an operant training preceded the motivational evaluation.

*Habituation*. For three consecutive days, animals could explore during 10 min the operant chambers, with an empty and lit food receptacle in one side, and two unilluminated identical holes in the opposite side. On the third day, they could find food in the food receptacle.

*Training*. After the habituation, mice were trained to nosepoke in one of the holes on a fixed ratio (FR)-1 schedule of reinforcement, where a single nose-poke elicited the delivery of a food pellet. The lit hole was designated as “active” and triggered the reward delivery.

Five seconds timeout to the FR-1 was given after a nose-poke, for both, right or wrong choices. During the timeout, additional nosepokes did not result in reward delivery. Each training session lasted an hour, or until a maximum of 50 rewards was delivered. The animals need to reach the following acquisition criteria before the schedule was increased to FR-5: (1) discrimination index of 3:1 for the active versus inactive responses, (2) obtaining 20 rewards per session (3) over three consecutive sessions. The FR-5 training lasted three consecutive days.

*Progressive Ratio Schedule of Reinforcement*. The ratio schedule for the PR testing was calculated as described by Richardson and Roberts (1996) using the following formula: $[5e^{(R \times 0.2)}] - 5$, where R is the number of rewards already earned plus 1. The ratios to obtain a food reward are 1, 2, 4, 6, 9, 12, 15, 20, 25, 32, 40, 50, 62, 77, 95, and so on. The last ratio completed by an animal is its breakpoint (BP). A PR session lasted an hour. The PR testing finished when the number of rewards earned in a session deviated less than 10% for three consecutive days.

**Additional Table 1.** ***Bregma coordinates and final size samples of the brain regions investigated***

| ROI | | *Bregma* coord. (mm) | Final n (ZO-1/IBA-1) | Final n (GFAP) |
|---|---|---|---|---|
| Prelimbic cortex | PL | 1.78–1.54 aB | 12 | 12 |
| Infralimbic cortex | IL | | 12 | 12 |
| Orbitofrontal cortex | OFC | | 12 | 11 |
| Nucleus Accumbens, core | NAcC | | 12 | 11 |
| Nucleus Accumbens, shell | NAcS | | 12 | 12 |
| Rostral anterior cingulate cortex | r-aCC | 1.10–0.86 aB | 12 | 12 |
| Dorsolateral striatum | DLS | | 12 | 12 |
| Dorsomedial striatum | DMS | | 12 | 12 |
| Olfactory tubercle | OTu | | 12 | 12 |
| Caudal anterior cingulate cortex | c-aCC | 0.26 aB–0.10 pB | 12 | 12 |
| **Anterior insular cortex** | aIC | | 12 | 12 |
| Paraventricular nucleus of the hypothalamus | PVN | | 12 | 12 |
| **Dorsal region of the bed nucleus of the stria terminalis** | dBNST | | 11 | 11 |
| Dorsal hippocampus | dHipp | 1.06–1.46 pB | 12 | 12 |
| Central amygdala | CeA | | 12 | 12 |
| **Basolateral amygdala** | BLA | | 12 | 12 |
| Rostral ventromedial hypothalamus | r-VMH | | 12 | 12 |
| Lateral habenula | LHb | 1.70–2.18 pB | 12 | 12 |
| Caudal ventromedial hypothalamus | c-VMH | | 12 | 12 |
| Subthalamic nucleus of the hypothalamus | STN | 2.92–3.16 pB | 11 | 12 |
| Ventral hippocampus | vHipp | | 10 | 10 |
| **Ventral tegmental area** | VTA | | 11 | 11 |

**Additional Figure 1. Relative expression of biomarkers of neurovascular components in a wide selection of brain regions.** ZO-1 expression for tight junctions of endothelial cells; IBA-1 expression for microglial cells; GFAP expression for astrocytes.

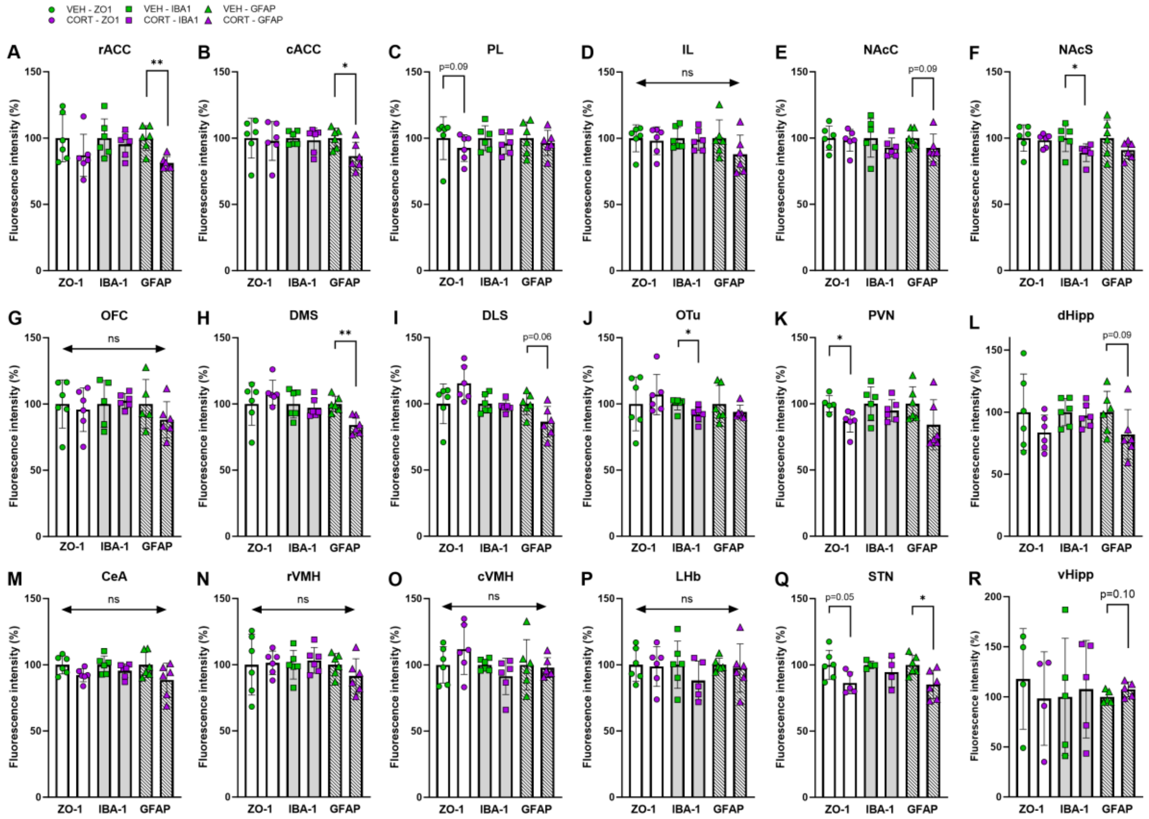

**Additional Figure 2. Surface occupied by biomarkers of neurovascular components in a wide selection of brain regions in control (VEH) and distressed (CORT) mice.** ZO-1 for tight junctions of endothelial cells; IBA-1 for microglial cells; GFAP for astrocytes.

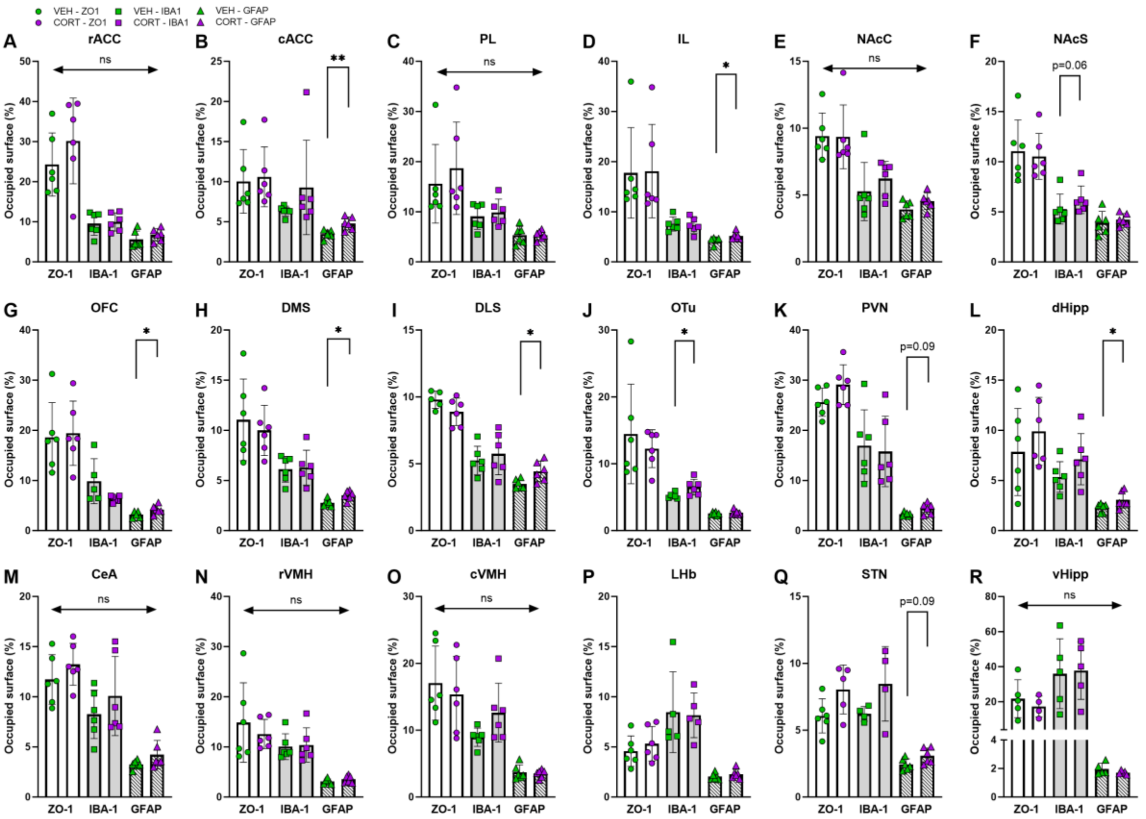


**Additional Figure 3. Linear correlations between ZO-1 relative expression and the surface occupied by it in 4 brain regions of interest** (aIC, dBNST, BLA and VTA) **in control** (green) **and distressed** (violet) **mice.**

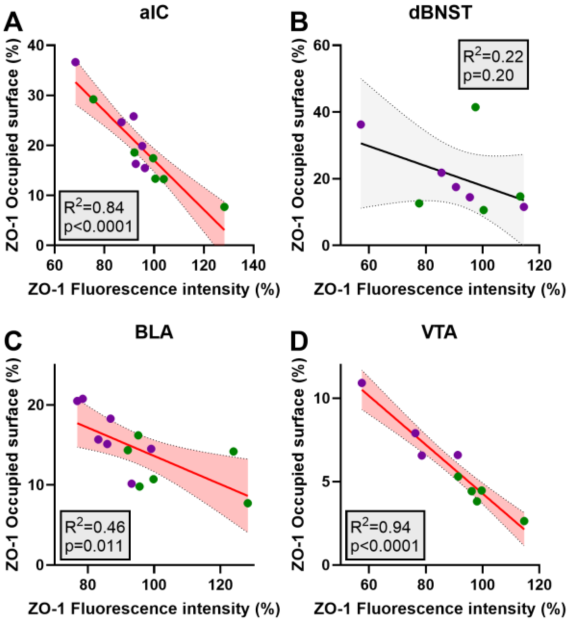

**Additional Figure 4. Microglial density** (number of cells per unit of surface) **in (A) the 4 main brain regions of interest for the present study, and (B) other wide number of brain regions, in control (VEH) and distressed (CORT) mice.**

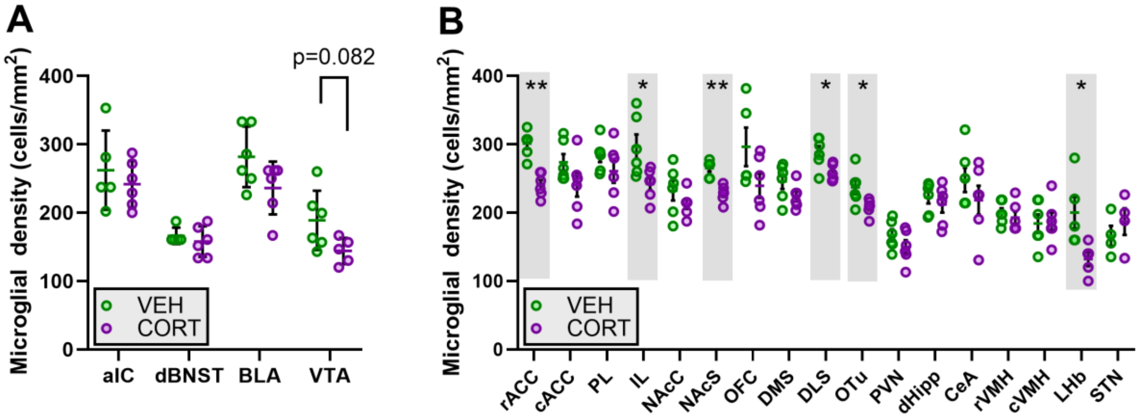


**Additional Figure 5. Linear correlations between motivational scores of control (VEH) and distressed (CORT) mice in the PR schedule of reinforcement task, and the measured NVU components.**

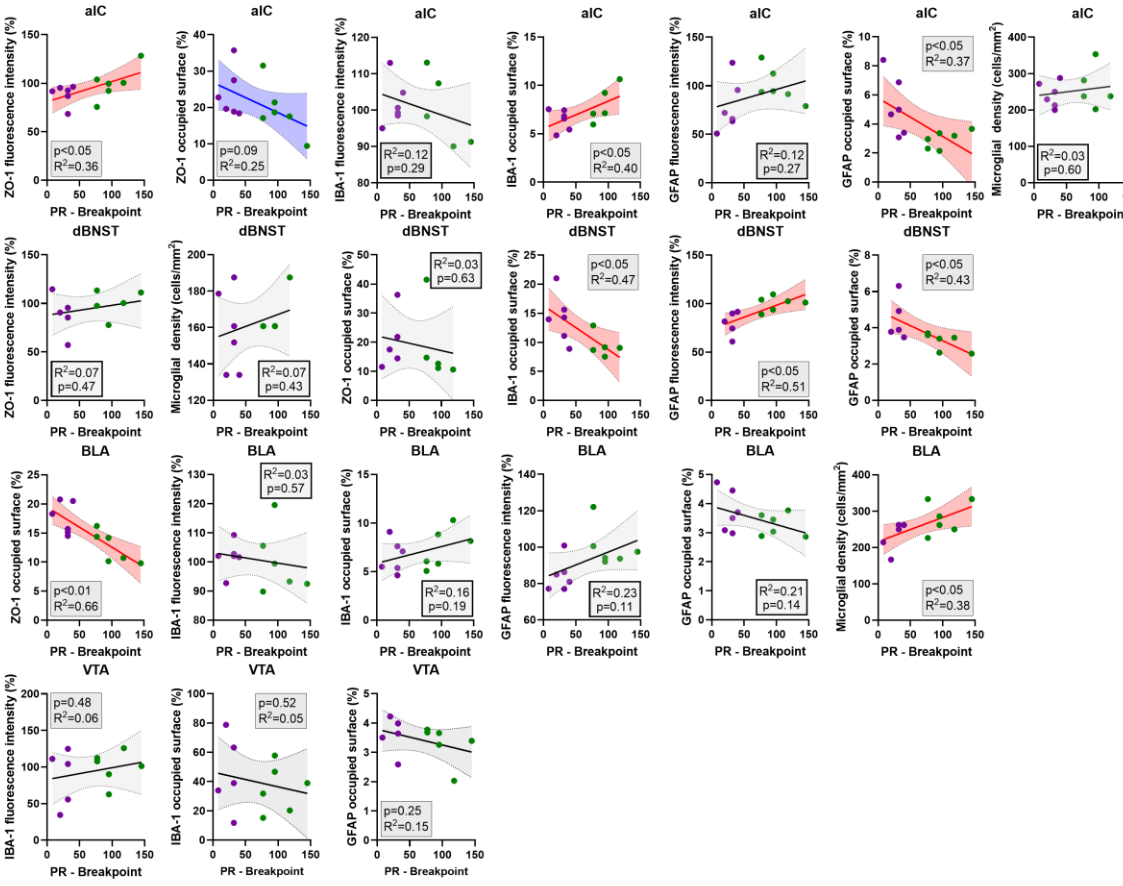

**Additional Figure 6. Brain interregional relationships between NVU components of control (VEH) and distressed (CORT) mice in the 4 main brain regions of interest for the present study.**

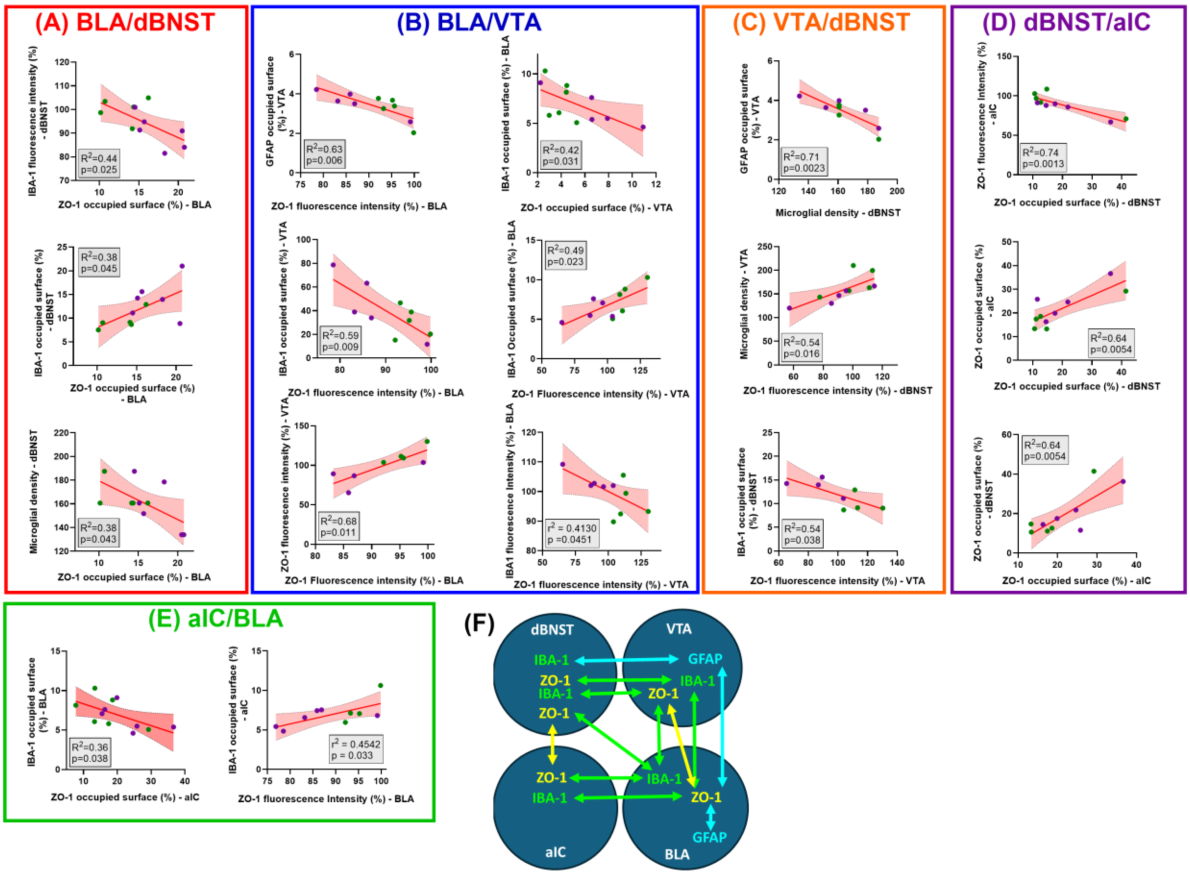
